# Loss of neuronal population organization links pathology to behavior in a model of Alzheimer’s disease

**DOI:** 10.64898/2026.03.18.712735

**Authors:** Douglas A. Ruff, Drew E.G. Sheets, Ramanujan Srinath, Giovanne B. Diniz, Devon J. Griggs, Danielle Beckman, Sean Ott, Kayla Schwartz, Carissa Erices, Scott Muller, Jeffrey H. Kordower, John H. Morrison, Marlene R. Cohen

## Abstract

Alzheimer’s disease (AD) and related dementias (ADRD) are defined by molecular and cellular pathology and cognitive decline, but linking these levels requires understanding how pathology alters large-scale neuronal activity. We longitudinally tracked behavior, multi-area neuronal population activity, and fluid and histological biomarkers in a macaque model of early-stage ADRD. As pathology progressed, visually guided behavior became increasingly disorganized, reflected in less structured exploration despite preserved task performance. Guided by systems neuroscience principles linking neuronal population activity with organized goal-directed behavior, we found progressive reductions in coordinated neuronal population activity within and between visual and parietal cortices, even as single-neuron tuning and basic feature encoding remained stable. These changes emerged when tau pathology was largely confined to regions providing feedback to visual cortex. This disorganized state appears modifiable: proof-of-concept methylphenidate administration was associated with transient improvement in behavioral organization. Together, these findings identify disruption of neuronal population organization as a defining feature of early-stage ADRD and frame early dysfunction as a disorder of coordinated population activity.

## Introduction

Alzheimer’s disease (AD) and related dementias (ADRD) gradually erode the brain’s ability to support flexible, goal-directed behavior, yet the earliest signs of dysfunction are often difficult to detect. Standard cognitive tests typically reveal impairments in executive function and declarative memory only after substantial neuronal loss has already occurred (Dubois et al., 2016; Jack et al., 2024; Jessen et al., 2014; Porsteinsson et al., 2021). Although considerable progress has been made in characterizing molecular pathology, the neural circuit mechanisms linking early pathology to cognitive dysfunction remain poorly understood (Canter et al., 2016; Grieco et al., 2023). In addition, many common animal models do not capture the subtle circuit-level disruptions or human-like behavioral changes that precede overt decline. These challenges limit the ability to identify mechanisms and to test interventions at a stage where they may be most effective (McKean et al., 2021). Addressing this problem requires approaches that link precise, quantitative measurements of behavior to the activity of neuronal populations across the brain.

Visually guided eye movements provide a powerful system for meeting this challenge. During natural vision, human and nonhuman primates typically make four to five eye movements per second, continuously integrating visual information, behavioral goals, and internal states (Ibbotson and Krekelberg, 2011; Krauzlis et al., 2017; Martinez-Conde et al., 2004; Sommer and Wurtz, 2008). These behaviors are rich but highly structured and quantifiable, and decades of basic neuroscience research, particularly in non-human primates, have established strong links between neuronal population activity and visually guided behavior (Parker and Newsome, 1998). Human studies suggest that eye movements are particularly sensitive probes of early AD deficits (Amieva et al., 2004; Molitor et al., 2015). Finally, the neural underpinnings of flexible exploration and goal-directed eye movements are well studied. They are supported by coordinated activity across multiple brain regions, including early and mid-level visual cortex and association areas involved in attention, decision-making, and working memory (Ebitz and Hayden, 2021; Ruff et al., 2018).

Here, we leverage this well-characterized system to link mechanistic insights and analytic approaches from systems neuroscience to translational questions by longitudinally measuring behavior, neurophysiology, and biomarkers in a new macaque model of ADRD (Beckman et al., 2021; Beckman et al., 2024). Rather than focusing on the loss of individual functions or representations, we examine how the disease progressively alters the organization of behavior and neuronal population activity. We had three goals: first, to identify early changes in the structure of visually guided behavior that emerge before overt impairments; second, to determine how these behavioral changes relate to degradation in the structure of neuronal population activity, despite preserved single-neuron tuning; and third, to define population-level signatures of circuit dysfunction that can guide targeted interventions. By linking subtle behavioral disorganization to disruptions in coordinated population activity, this work demonstrates that early disease progression is best characterized as a disorder of organization spanning scales of analysis from molecules to neuronal populations and behavior.

## Results

### Adeno-associated virus injection induces a macaque model of AD

The entorhinal cortex (ERC) is among the earliest and most severely affected regions in most human AD patients (Igarashi, 2023). In our macaque model, pathology is initiated by injecting an adeno-associated virus (AAV) expressing a double human tau mutation (P301L/S320F) into the ERC (Fig 1A). This manipulation produces local tau accumulation and later cell death that spreads gradually to connected regions, with a progression pattern and anatomical specificity that closely resemble human ADRD. The speed of progression is inversely related to the synaptic distance from the ERC, consistent with a spread of pathology that follows corticocortical and hippocampal connectivity (Beckman *et al*., 2021; Beckman *et al*., 2024).

**Figure 1.**
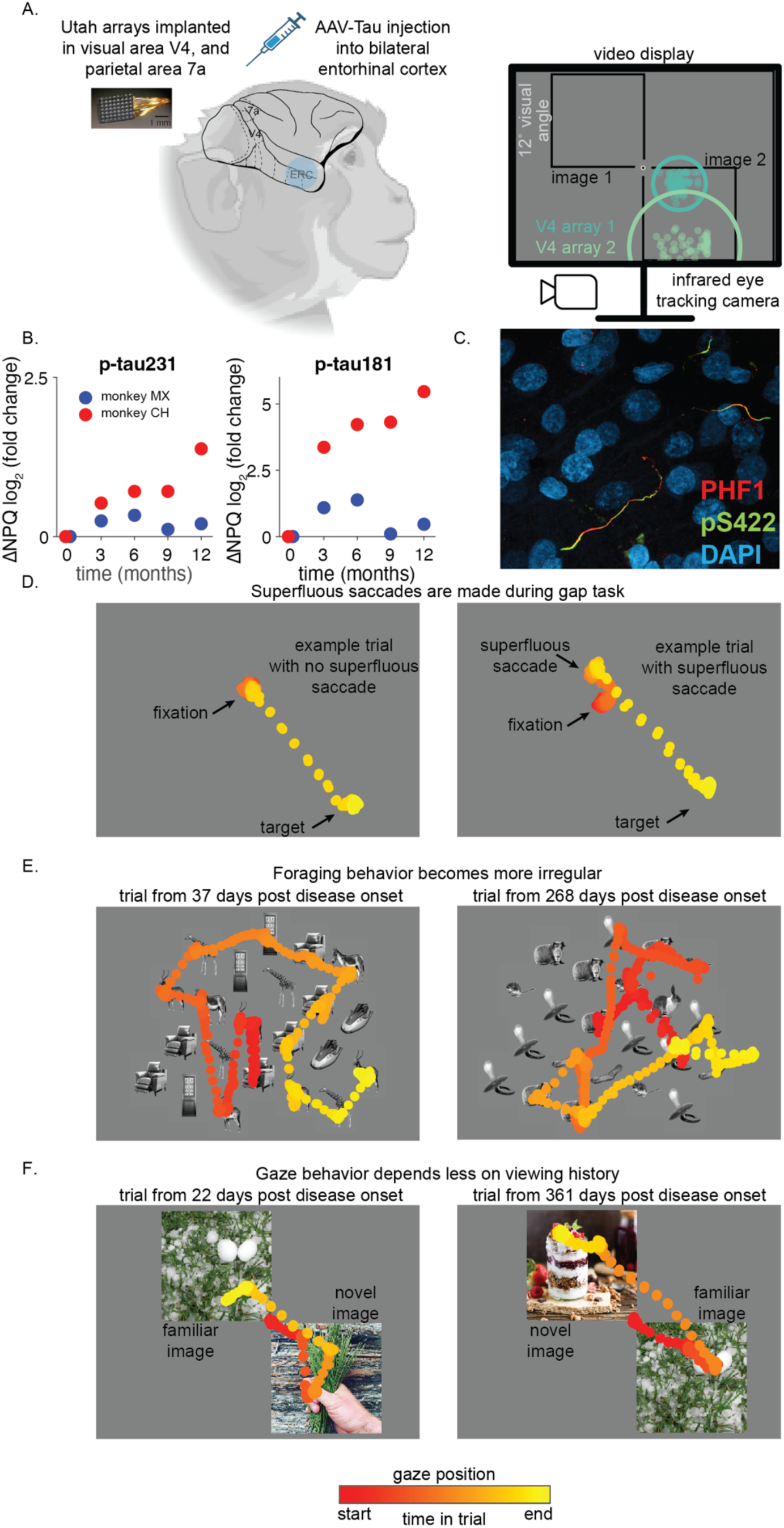
Experimental design and behavioral observations. A) Two monkeys received bilateral AAV-Tau injections into entorhinal cortex to induce progressive tauopathy (see Methods; (Beckman *et al*., 2021; Beckman *et al*., 2024)). Three 64-channel microelectrode arrays were chronically implanted: two in visual area V4 and one parietal area 7a. Estimates of the centers of the spatial receptive fields of units from the two V4 arrays relative to fixation from monkey CH are drawn as points on the computer monitor along with an outline of the visual stimulus configuration during the preferential looking task (described below). An estimate of the size of one unit’s receptive field from each array is drawn as a circle. Monkeys performed visually guided tasks based on images presented on a computer screen, controlled by infrared eye tracking, for one year following the injections while we simultaneously recorded neuronal responses. B) AAV-induced changes in plasma markers associated with early stages of ADRD were observed in both animals (data are plotted in units of log2 -fold change (Feng et al., 2023)) and C) histological signs of phosphorylated tau (PHF1, red; pS422, green; overlap, yellow; nuclei, DAPI, blue) were observed in V4. Sparse labeling of phosphorylated tau was seen in visual and parietal cortex and was predominantly restricted to processes, suggesting that functional changes observed in V4 and 7a do not arise primarily from tau-related cell death in those areas, and implicate synaptic and/or network mechanisms (image from right hemisphere of monkey CH shown, see Supplementary Figure 2). D) Two example trials from a gap task where a fixation spot turned off prior to the appearance of a saccade target. On most trials (left), monkeys waited to move their eyes until the appearance of a peripheral target and then made a single saccade to that target. On some trials (right), monkeys made an unnecessary and unrewarded saccade (called a superfluous saccade) prior to the appearance of the peripheral target. E) Two example scan paths during a visual foraging task where fixating certain stimuli (i.e. animals) was rewarded while fixations to others (i.e. objects) were unrewarded. The scan path from the example trial on the right appears more irregular, with larger distances between consecutive fixations and unrewarded returns to previously viewed locations. F) Scan paths from two example trials during a free-viewing preferential looking task. Monkeys were rewarded after accumulating one second of total view time on either of the images. In the trial on the left, the monkey looked first and longer at the novel image, explored it, and then switched once to look at the familiar image. On the right, the monkey looked first at the familiar image, and rapidly switched to the novel image. Monkey image in A created using BioRender.

Over the course of disease progression, we measured biomarkers, behavior, and neuronal population responses to obtain a comprehensive picture of the biological, neurophysiological, and behavioral changes. We used blood and cerebrospinal fluid measurements of phosphorylated tau, which are sensitive tools for detecting and staging AD in humans (Dubois *et al*., 2016; Hampel et al., 2004), to compare to human disease. In particular, plasma p-tau231 is thought to reflect early disease-related changes (Milà-Alomà et al., 2022), and p-tau181 has been shown to predict cognitive decline (Abdelnour et al., 2024; Tropea et al., 2023). In our monkeys, both biomarkers evolved as previously reported, confirming induction and spread of tau pathology and supporting the clinical relevance of this macaque model (Figure 1B; (Beckman *et al*., 2021; Beckman *et al*., 2024)).

We performed histological analyses one year after disease onset to assess the extent and distribution of pathology in brain areas relevant to this study. In both animals, there were clear signs of abnormally hyperphosphorylated tau throughout the medial temporal lobe (MTL; Supplementary Figure 1), consistent with previous reports using this model (Beckman *et al*., 2021; Beckman *et al*., 2024). Tau pathology outside the MTL was detectable but remained limited. In area V4, where the majority of physiological recordings in this study were obtained, tau pathology was observed predominantly in processes, likely belonging to feedback projections (Figure 1C, Supplementary Figure 2). Together, these findings indicate that the disease-related changes observed in V4 should not be interpreted as reflecting lesions at the site of our electrodes. Instead, they are more consistent with alterations to the broader network influencing V4 activity and the visually guided behaviors examined in this study. These results are consistent with our monkeys reaching Braak stage 2-3 classification, which is thought to correspond to early to mid-stages of the disease (Braak et al., 2006; Braak and Braak, 1991). Our results therefore highlight changes that occur during the earliest stages of disease progression.

### Early behavioral disorganization precedes overt functional decline

The longitudinal design of our study allowed each animal to serve as its own control, providing high sensitivity to detect subtle, progressive changes (Jessen *et al*., 2014). We trained behavioral tasks before injection of the disease construct and recorded behavioral and neuronal data in daily sessions for one year post disease onset. Many aspects of behavior and neuronal responses remained stable: monkeys performed trained tasks effectively, and social, feeding, and general behaviors in the home environment were largely unchanged.

We evaluated visually guided eye movement behaviors in three tasks designed to probe cognitive processes that we hypothesized would change early in disease progression. Despite overall stable performance, we observed rapid, subtle, and consistent changes in how visually guided behaviors were organized.

- Directed saccade (“gap”) task: Early AD in humans is associated with diminished ability to suppress anticipatory or inappropriate saccades, suggesting impaired behavioral inhibition (Abel et al., 2002). To test for similar deficits, we used a classical “gap task” that manipulates the relative timing between the disappearance of a fixation spot and the appearance of a peripheral target (Abel *et al*., 2002). As disease advanced, monkeys became more likely to make unrewarded superfluous eye movements before target onset (Figure 1D). Notably, they continued to generate accurate saccades to targets, indicating a selective decline in inhibitory control rather than a general impairment of oculomotor function.
- Rule-based visual foraging task: Human AD patients show impairments in planning and organizing visual search behavior (Molitor *et al*., 2015; Moser et al., 1995; Rösler et al., 2000). We assessed these processes using a rule-based foraging task where monkeys were rewarded for fixating only some target images (e.g. animals) and not distractors (e.g. inanimate objects). As disease progressed, monkeys sampled the display more irregularly, consistent with a breakdown in the organization of goal-directed exploration (Figure 1E).
- Preferential looking task: Humans with AD and mild-cognitive impairment and animal models with hippocampal damage have altered gaze patterns during free viewing, particularly showing reduced preference for novel compared to familiar images (Bott et al., 2017; Crutcher et al., 2009; Lagun et al., 2011; Zola et al., 2013; Zola et al., 2000). In a preferential looking task inspired by these studies, monkeys displayed reduced preference for fixating novel relative to familiar images and switched their gaze between images more frequently (Figure 1F). These changes parallel those reported early in human AD and in monkeys with hippocampal lesions (Bastin et al., 2019; Bott *et al*., 2017; Buffalo et al., 2000; Chau et al., 2017; Crutcher *et al*., 2009; Daffner et al., 1992; Lagun *et al*., 2011; Ryan et al., 2000; Zola *et al*., 2013; Zola *et al*., 2000).

Example trials (Figure 1) suggest that behavior becomes less organized, less strategic, and faster during disease progression. We quantified these observations as a function of time since disease onset, defining day 0 as the day of AAV-tau injection in entorhinal cortex. Although overall task performance remained stable, multiple indicators of behavioral organization changed rapidly and systematically.

In the directed saccade task, monkeys continued to generate accurate saccades to peripheral targets. However, they increasingly initiated premature (superfluous) saccades after the fixation point was extinguished but before the appearance of the target (Figure 2A), suggesting a progressive decline in behavioral inhibition, without evidence of more general saccadic impairment.

**Figure 2.**
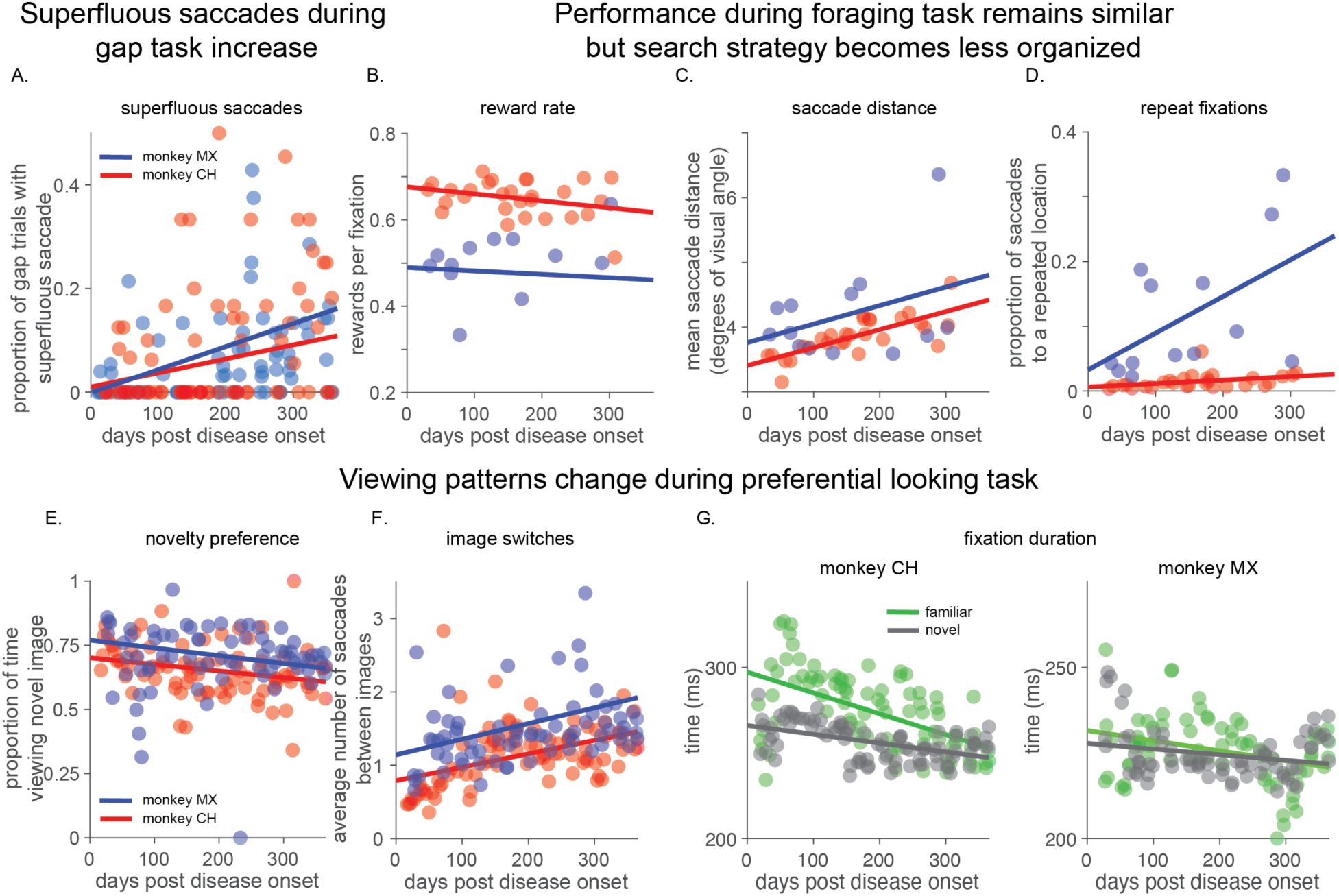
Progressive disorganization of visually guided behavior. A) The frequency of trials in the gap task in which monkeys made an unrewarded superfluous saccade before the target appeared increased across one year post disease onset (both slopes p < 0.001, all slope tests are F-tests on a linear model against the null hypothesis of a constant slope) B)-D) Performance during visual foraging task. B) Reward rate per fixation remained largely stable (monkey CH, p = 0.12; monkey MX, p = 0.8). C) The mean distance between consecutive saccade endpoints increased in one animal, with a similar trend in the other (monkey CH, p < 0.001; monkey MX, p = 0.21). D) The rate of revisiting a location during a trial, which was always unrewarded, increased (monkey CH p < 0.05; monkey MX p < 0.05). E)-G) Free viewing during the preferential looking task. E) Preference for viewing novel images decreased (monkey CH, p < 0.001; monkey MX, p < 0.05). F) The number of saccades that switch between images during a trial increased (monkey CH, p < 0.001; monkey MX, p < 0.05). G) The amount of time per fixation on both novel and familiar images decreased, each monkey plotted separately (monkey CH, left, familiar p << 0.001, novel p << 0.001; monkey MX, right, familiar p < 0.005, novel p < 0.05). Together, these results indicate a gradual loss of structured, goal-directed exploration and reflect early behavioral disorganization.

In the rule-based visual foraging task, monkeys continued to distinguish targets from distractors, and their rates of selecting targets remained relatively stable (Figure 2B). However, their exploration became increasingly disorganized. The average distance between consecutive fixations gradually increased, as did the frequency of revisiting previously sampled locations (which was never rewarded). These metrics indicate increasing variability and reduced structure in goal-directed exploration (Figure 2C, D).

In the preferential looking task, overall engagement with the stimuli remained relatively stable, but viewing patterns changed markedly. The established preference for novel over familiar images declined gradually (Figure 2E). The monkeys also switched their gaze between the two images more frequently (Figure 2F), and the duration of fixations on novel and familiar images became shorter (Figure 2G; (Bakotich et al., 2024; Hannula et al., 2010; Smith et al., 2006; Smith and Squire, 2008)). These changes are consistent with faster, less strategic, and less organized visual exploration.

Together, these results reveal that early ADRD is not defined by failure, but by a progressive degradation of the structure of visually guided behavior. These changes are not easily detectable through coarse or unconstrained behavioral observation but are readily apparent through sensitive, longitudinal measurements.

### Stimulus representations remain intact while population structure deteriorates

The entorhinal cortex, the site of our AAV-injection, is strongly interconnected with many brain areas, including the ventral visual stream and association areas (Garcia and Buffalo, 2020). Given converging evidence for early visual cognitive impairments in human AD (Hannula *et al*., 2010; Molitor *et al*., 2015; Opwonya et al., 2022) and the role of areas like mid-level visual V4 and parietal area 7a in the processes impacted in our behavioral tasks (including visually guided exploration, attention, and visual memory (DiRisio et al., 2025; Motley et al., 2018; Rozzi et al., 2006; Ruff *et al*., 2018)), we asked whether early disease progression alters how visual information is represented in those neuronal populations.

Because behavior remained guided by visual information but became disorganized, we hypothesized that neurons in visual cortex would continue to encode visual features, but that the organization of responses across neuronal populations would degrade during disease progression. To test this, we recorded multi-unit responses of populations of neurons in area V4 to a range of visual stimuli at various sizes and positions, presented within and outside their classical receptive fields.

Early in disease progression, neuronal populations in V4 exhibited canonical organization for basic features. As in healthy animals, units were tuned for the expected mid-level visual properties (Figure 3A; (Pasupathy et al., 2020; Roe et al., 2012) and stimulus features that are not typically correlated in natural scenes, such as color and shape, were encoded independently and robustly in the population (Srinath et al., 2024) (Figure 3B). Over time, despite the continued presence of tuned units (Figure 3C), this independence broke down and population responses to different stimuli appeared to become less discriminable (Figure 3D).

**Figure 3.**
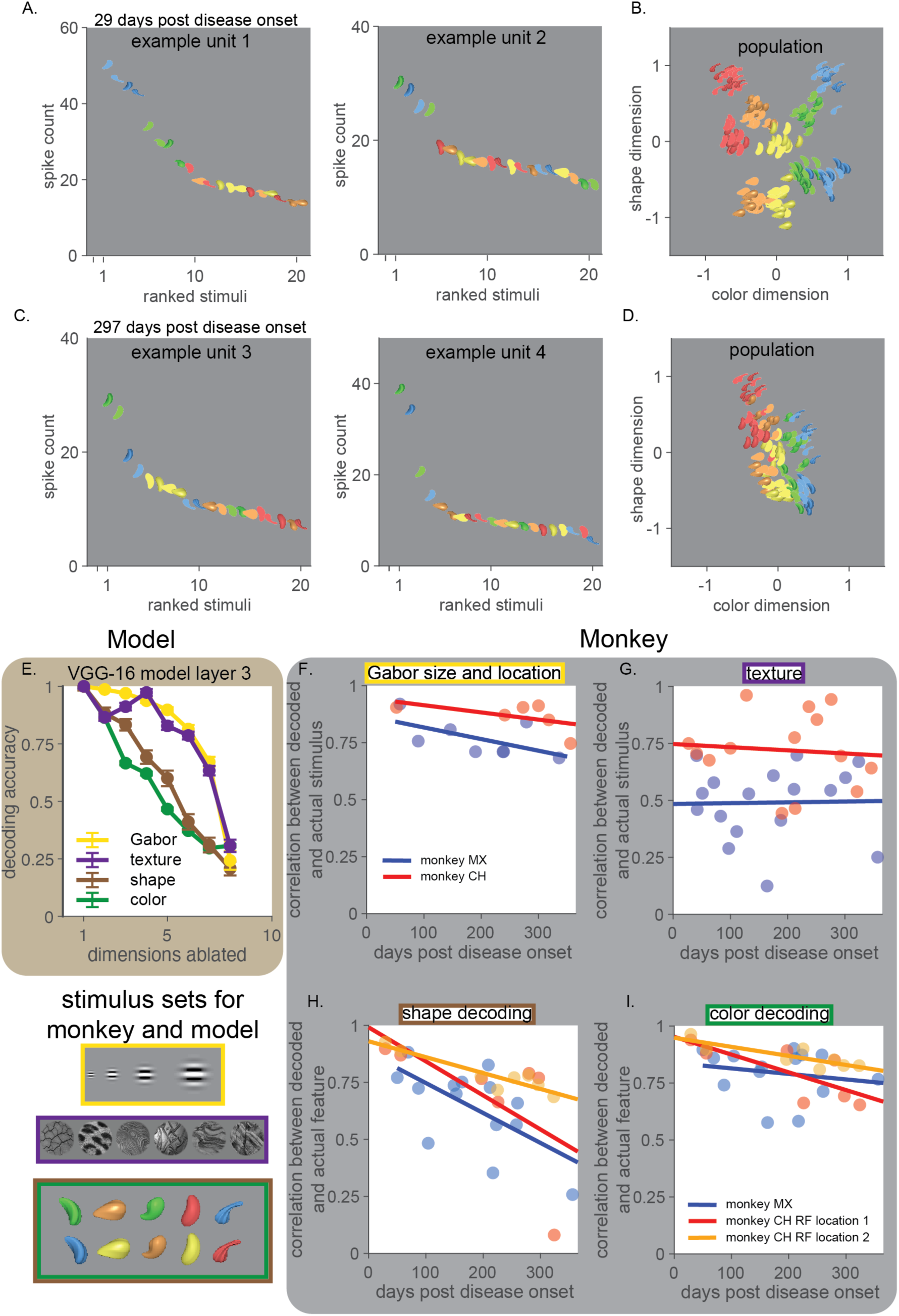
Population-level visual representations lose organization despite stable responses to many visual features. A) Example responses to a set of colored shape stimuli (Srinath *et al*., 2024) from two example V4 units 29 days post disease onset. Visual stimuli are represented by icons and ordered on the x-axis by average response magnitude during a 200ms window, shifted 50ms to account for response latency (y-axis). B) Projections of V4 population responses from the same example session 29 days post disease onset onto axes in neuronal population space that account for the most color and shape information (identified using QR orthogonalization; see Methods). Each icon represents the response of the population to that stimulus on a single trial. C)-D) Same as A and B from an experimental session 297 days post onset. While tuning for the stimulus features remained, population responses to different stimuli appear less separable. E) A convolutional neural network model (VGG-16) was used to generate hypotheses about how the disruption of local computations might alter the information about different stimulus features encoded in V4 populations. Removing dimensions of population activity in a V4-like layer of the model more precipitously affected the information represented about our colored shape images compared to texture and Gabor stimuli. F) As predicted by the model, the ability to linearly decode the size and location of a Gabor stimulus from the responses of the recorded V4 neurons remained unchanged for one year post disease onset (see Methods, monkey CH, p = 0.27, monkey MX p =0.13, all slope tests are F-tests on a linear model against the null hypothesis of a constant slope). G) Similarly, the ability to distinguish between different visual texture stimuli remained unchanged (monkey CH, p = 0.76, monkey MX, p = 0.93). In contrast, the ability to linearly decode H) shape and I) color information declined post disease onset (two stimulus locations for monkey CH that overlap two clusters of receptive fields, see Methods, shape: monkey CH p < 0.0001 & p = 0.13; monkey MX, p < 0.01; color: monkey CH: p < 0.001 & p < 0.05; monkey MX, p = 0.51).

The anatomical spread of tau from late to early ventral stream areas that was observed in previous work (Beckman *et al*., 2021; Beckman *et al*., 2024) led us to hypothesize that the representations of features that rely on computations within later or middle ventral areas would degrade earlier than representations of features that rely only on early stages. To assess which features of our stimuli rely on later stages, we selectively removed individual dimensions of the activity of a convolutional neural network (CNN, see Methods). CNNs model hierarchical visual processing in the ventral visual stream (Lindsay, 2021; Yamins and DiCarlo, 2016) and allow us to perform ‘ablation’ experiments to assess the impact of different stages of processing on the way different stimulus features are encoded.

Randomly removing dimensions of population activity from a V4-like layer of the model disrupted the encoding of some visual features more than others (Figure 3E). The representation of simple stimulus features (like the spectral components of Gabors, or the discrimination of textures), which are likely inherited from earlier layers, were more robust to ablations in the activity of a V4-like layer than were representations of the shape and color of more complex images (Figure 3E).

This prediction was confirmed in our V4 population recordings. The ability to linearly decode low-level visual features like the position and size of Gabor stimuli or the identity of different visual textures remained relatively stable (Figure 3F and G). This stability also indicates that gradual changes in recording quality did not necessitate a decline in the information available in the recorded population. In contrast, information about the shape and color of complex stimuli declined over time (Figure 3H and I), consistent with a selective degradation of the population structure that supports coordinated encoding of complex features.

Together, these results indicate that early in disease progression, visual feature encoding, especially for features that could be inherited from earlier ventral areas, remains largely intact, even as the structure of population responses degrades. This dissociation reveals that early pathology spares local feature tuning while disrupting signals that organize and coordinate neuronal populations.

### Early disruption of signals that organize neuronal populations

We therefore asked whether population signatures known to reflect coordinated processing across neurons are disrupted early in disease. Previous work, including from our lab, has demonstrated that the structure of population activity in visual cortex reflects not only stimulus-driven tuning and patterns of connectivity, but also internal cognitive states like attention, arousal, and task uncertainty (Ruff *et al*., 2018). These internal states influence the structure and extent to which neuronal responses co-fluctuate from trial-to-trial. This covariability is often quantified using pairwise noise correlations (also called spike count correlations or r_SC_, (Cohen and Kohn, 2011)) and by analyzing the geometry of population responses to repeated presentations of the same stimuli (Azeredo da Silveira and Rieke, 2021; Chung and Abbott, 2021; Kriegeskorte and Wei, 2021). These population-level signatures are widely interpreted as consequences of local and long-range connectivity, including feedback signals from downstream areas, that shape and organize population responses (Semedo et al., 2022; Smith et al., 2013).

In this monkey model, tau pathology was largely confined to medial temporal and connected regions, with little direct accumulation or damage to V4 and 7a at the stages we examined (Beckman *et al*., 2021; Beckman *et al*., 2024). We therefore hypothesized that V4 and 7a population activity would reflect disruption of network effects like feedback and inter-area coordination, which could be measurable as disruption of correlated variability consistent with a decline in coordination of population activity. We tested this hypothesis by examining population activity in V4, and 7a during the brief fixation period at the start of each trial, when the screen was blank and stimulus-driven responses were minimal.

Consistent with our hypothesis, we observed an early and progressive decline in these population signatures. During disease progression, correlated variability decreased both within each area and between V4 and 7a (Figure 4A-C). In parallel, estimates of intrinsic timescale, which quantify how strongly past neuronal activity influences current response dynamics, also declined (Figure 4D; (Soyuhos et al., 2025)).

**Figure 4.**
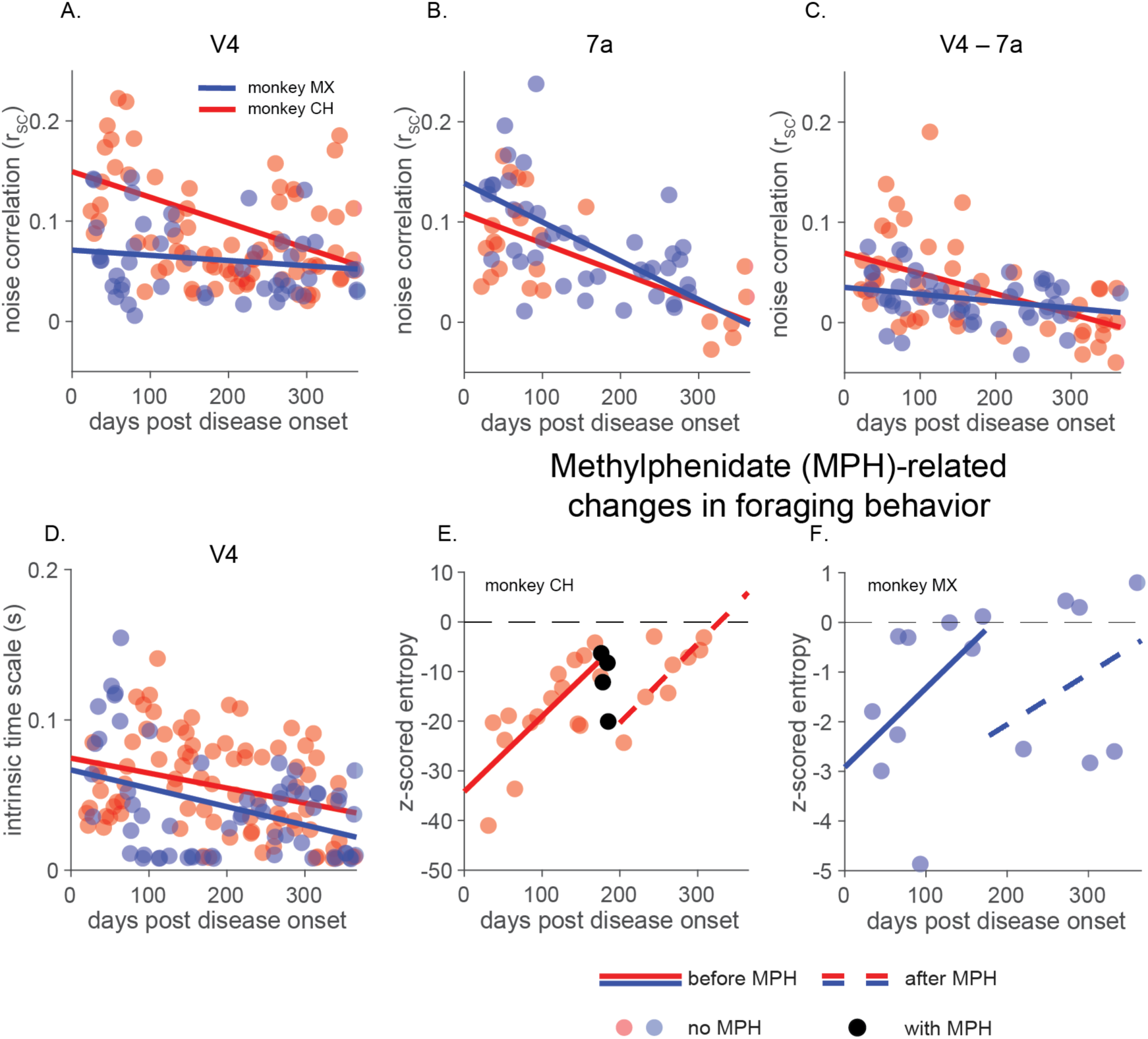
Early disruption of feedback-associated population signals and proof-of-concept intervention with methylphenidate (MPH). A-C) Shared variability decreased during disease progression. The plots show firing-rate matched noise correlations (r_SC,_; see Methods) averaged across simultaneously recorded pairs of units measured during a 200ms period of stable fixation before stimulus presentation. A) Mean shared variability from both monkeys decreased in V4 (monkey CH, p < 0.001, monkey MX p = 0.38, all slope tests are F-tests on a linear model against the null hypothesis of a constant slope) B) 7a (monkey CH, p < 0.01, monkey MX p < 0.001) and C) between V4 and 7a neurons (monkey CH, p < 0.001, monkey MX p < 0.05). D) The decay time of V4 autocorrelations (average intrinsic time scale) decreased throughout disease progression, suggesting altered circuit properties (see Methods; monkey CH, p < 0.005, monkey MX, p < 0.001). E) and F) Methylphenidate (administered orally) had long lasting effects on behavioral strategy (quantified as z-scored foraging entropy; see Methods) during the visual foraging task (slopes of best fit are not distinguishable pre vs post MPH administration; monkey CH p = 0.92; monkey MX p = 0.78).

Although the relationships between firing rate, correlation structure, and dimensionality are complex and nonlinear (Cunningham and Yu, 2014; Panzeri et al., 2022; Tian et al., 2025; Umakantha et al., 2021), the pattern we observe indicates a progressive reduction in coordinated population activity. Neuronal firing rates fluctuate more independently of one another, consistent with a gradual disruption of feedback from higher-order areas and/or recurrent interactions within each area. These findings link the preserved stimulus encoding to the early behavioral disorganization we observed in this model of ADRD.

### Proof of concept: using neuronal population activity to guide targeted interventions

The results above reveal a progressive loss of organization in neuronal population activity, even as representations of visual features likely inherited from earlier stages of the ventral visual stream remain relatively preserved. This dissociation indicates that early disease progression disrupts the coordination of neuronal responses before it produces broad degradation of sensory representations. These population-level changes parallel the subtle but consistent disorganization observed in visually guided behavior.

Previous systems neuroscience studies show that correlated variability is modulated by cognitive processes like attention, arousal, or task engagement (Ni et al., 2018; Ruff and Cohen, 2014; Ruff and Cohen, 2019; Ruff *et al*., 2018). We have also previously shown that methylphenidate (Ritalin), a psychostimulant commonly prescribed for attention deficit hyperactivity disorder, enhances visually guided behavior and modulates correlated variability in visual cortex (Ni et al., 2022). Some studies suggest that methylphenidate can reduce apathy in humans with AD (Drye et al., 2013; Kishi et al., 2020; Mintzer et al., 2021; Padala et al., 2017; Rosenberg et al., 2013; van Dyck et al., 2021), but its effects on the organization of visually guided behaviors are largely unknown. On this basis, we asked whether the disorganized state we observed during disease progression is modifiable using the same methods that change neuronal population signatures of organization in healthy animals.

To test this possibility, we administered methylphenidate (3.5 mg/kg, in the range we used in our previous study; (Ni *et al*., 2022)) beginning approximately six months after disease onset, when behavioral disorganization was measurable in both animals (see Methods). In monkey CH, the drug was administered across 18 days during three-plus weeks of testing starting on day 176, while in monkey MX it was given across 5 days during one week of testing starting on day 174. Monkey CH performed visually guided tasks during experimental sessions that immediately followed drug administration, while monkey MX declined to perform any task.

In both animals, methylphenidate partially restored organized visual foraging. Before treatment, the entropy of visual search patterns increased steadily over the first six months, reflecting a progressive loss of organized exploration. Following the onset of methylphenidate administration, entropy decreased sharply, returning to levels observed several months earlier. Once we stopped treatment, entropy did not return immediately to pretreatment levels, even though the direct impacts of methylphenidate should have dissipated within one day (Doerge et al., 2000; Ni *et al*., 2022; Oemisch et al., 2016). Instead, entropy began to slowly increase at a rate similar to pre-treatment (Figure 4E, F).

Although this preliminary intervention was not designed as a systematic evaluation of therapeutic efficacy, it demonstrates two key points. First, pharmacological interventions can transiently restore organized behavior even in the presence of ongoing pathology. Second, knowledge about how neuronal population activity organizes behavior can guide the identification and testing of candidate interventions that target circuit-level disruptions in cognitive disorders.

## Discussion

We show that early-stage pathology in a model of ADRD is characterized by disruption of neuronal population organization and parallel disorganization of behavior, even as single-neuron tuning and basic sensory encoding remain largely intact. These changes arose at stages when tau pathology in visual cortex remained limited, indicating that functional disruption extends beyond areas of prominent pathology. Coordinated population activity within and between visual and parietal cortices progressively declined as pathology advanced, and these changes were reflected in structured alterations in visually guided behavior. Together, these findings identify disruption of neuronal population organization as a defining feature of early-stage ADRD and suggest that early dysfunction reflects altered coordination across distributed cortical networks rather than degradation of sensory encoding.

Coordinated neuronal population activity organizes how information is combined across brain areas to guide behavior. When neurons fluctuate together in structured ways and cortical areas interact coherently, distributed representations can be integrated into stable, goal-directed actions rather than fragmented or inconsistent responses. The progressive reduction in coordinated population activity observed here therefore provides a systems-level explanation for why behavior becomes disorganized despite preserved single-neuron tuning (Chockanathan and Padmanabhan, 2022; Ebitz and Hayden, 2021; McGregor et al., 2024; Moghaddam and Wood, 2014). Early-stage pathology does not primarily eliminate sensory information; instead, it appears to disrupt the coordinated interactions that transform intact representations into organized behavior.

Early-stage ADRD has traditionally been conceptualized in terms of accumulating molecular pathology and progressive neuronal degeneration. While these processes are central to disease progression, established links between pathology and the cognitively complex behaviors affected in ADRD remain limited. A substantial body of work in systems neuroscience shows that coordinated neuronal population activity provides the substrate for organized, goal-directed behavior. Guided by these principles, we examined whether disruption of population organization could account for early behavioral changes in ADRD. By demonstrating that coordinated population activity declines early in this primate model, even when tau pathology in visual cortex remains limited and single-neuron encoding is preserved, we provide a causal test of these systems-level ideas in a disease context. These results further show that disorganization across levels of spatial scale, from molecular to neuronal populations, is an early feature of ADRD (Figure 5). Neuronal population organization thus emerges as a central characteristic of early-stage ADRD that remains, at least in part, modifiable.

**Figure 5.**
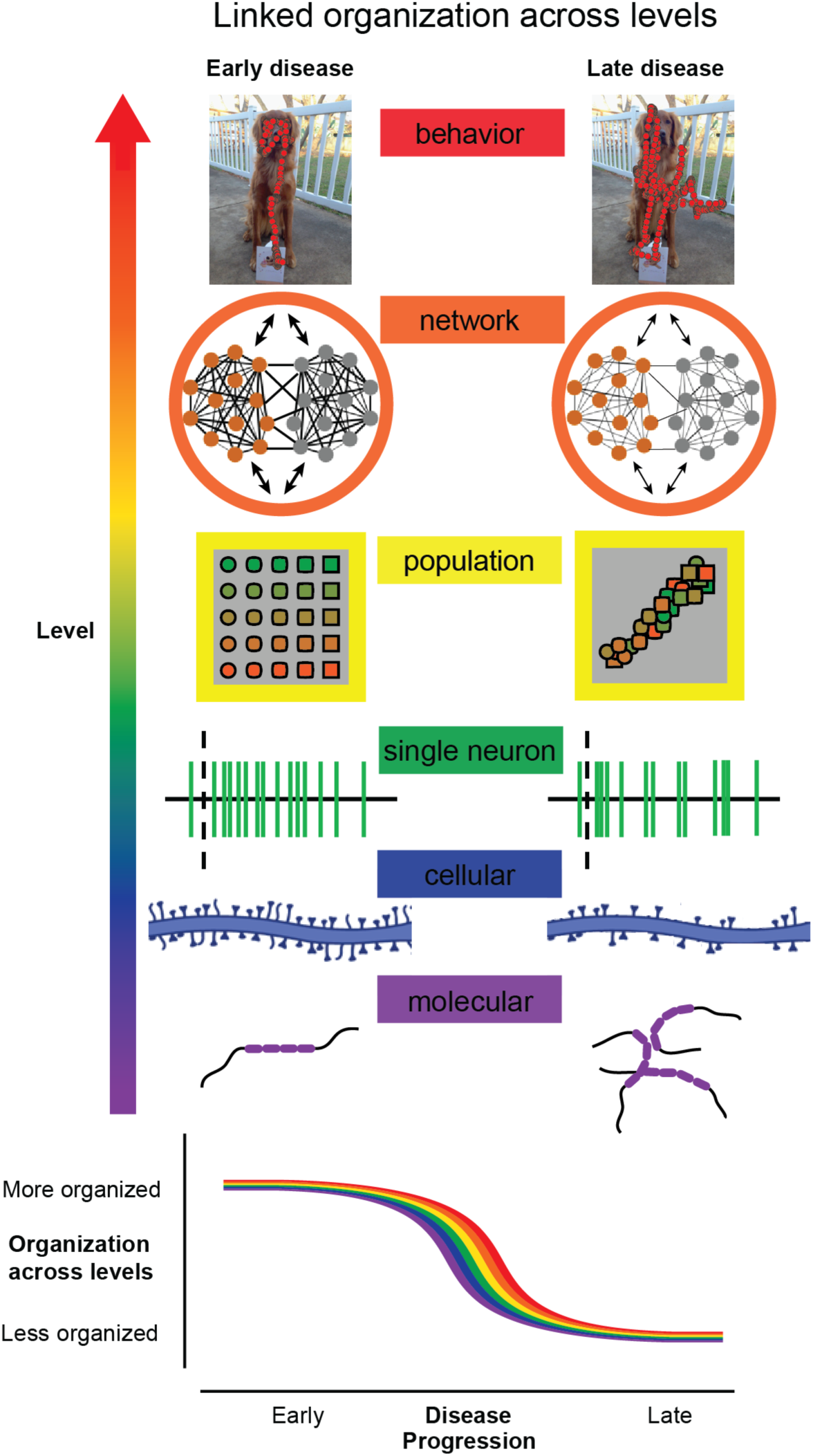
Loss of organization across levels links pathology to behavior. Organized behavior depends on coordinated structure across levels of the brain, from molecules to networks. In the healthy state, molecular organization (tau protein folding), cellular organization (intact synaptic connectivity), single-neuron activity (structured variability and intrinsic dynamics), population representations (separable feature encoding), and network interactions (coordinated activity within and across areas) together support goal-directed behavior (low entropy, structured, efficient). During disease progression, organization at each of these levels is disrupted: tau misfolding and accumulation, synaptic alterations, changes in intrinsic dynamics, reduced separability of population responses, and weakened inter-areal coordination. These changes emerge in parallel with increasing behavioral disorganization (higher entropy, less structured exploration). This schematic illustrates that disease progression reflects a progressive loss of coordinated organization across levels rather than a uniform loss of function. Molecular and cellular images were created using BioRender.

### Disorganization of coordinated population activity underlies early cognitive dysfunction

The earliest changes we observed during disease progression were not impairments in basic task performance or worsening of single-neuron tuning, but alterations in behavioral structure and strategy. Early in disease progression, monkeys continued to perform the basic visual tasks, but their behavior became progressively less structured, less strategic, and faster. These behavioral changes were mirrored at the population level: while representations of visual features likely inherited from earlier ventral stream areas remained relatively preserved, the coordination of neuronal responses deteriorated. Population signatures that, in healthy animals are modulated by attention, arousal, or task uncertainty (Maunsell, 2015; Ni *et al*., 2018; Ruff and Cohen, 2014; Ruff and Cohen, 2019; Ruff *et al*., 2018; Xue et al., 2024) declined early in disease progression, suggesting that cognitive symptoms may emerge from disruptions of the same network computations that support flexible behavior in healthy individuals.

By directly measuring changes in the activity of neuronal populations and quantifying the magnitude and structure of correlated variability, we show that the organization of population activity is altered early in disease progression in ways that are not readily apparent from single-neuron responses or coarse behavioral observation. Because these signatures are thought to depend on feedback and corticocortical interactions that support flexible behavior, their early disruption implicates those mechanisms and is consistent with the notion that corticocortical circuits interconnecting association regions are highly vulnerable in ADRD (Hof and Morrison, 1994).

### The degradation of neuronal population organization is partially modifiable

Because we observed that coordinated behavior and the coordinated neuronal population activity that underlies it deteriorates during disease progression, we asked whether this degraded state could be rescued. Previous work in systems neuroscience demonstrates that coordinated neuronal population activity is dynamically modulated by cognitive processes like attention, arousal, and learning (Maunsell, 2015; Ni *et al*., 2018; Ruff and Cohen, 2014; Ruff and Cohen, 2019; Ruff *et al*., 2018; Xue *et al*., 2024). We previously showed that methylphenidate can alter coordinated population activity and behavior in ways that resemble successful attention and arousal in healthy animals (Ni *et al*., 2022). On this basis of this foundation, we conducted a preliminary test of whether the disorganization arising during neurodegeneration could be partially reversed.

We found that methylphenidate temporarily restores behavioral organization in this model of early-stage ADRD. Although this preliminary demonstration was not designed as a systematic assessment of pharmacological efficacy, it indicates that the degradation of organized behavior and its neural underpinnings remain partially modifiable, at least in the early stages. These findings illustrate the potential for using neuronal population measurements and knowledge derived from systems neuroscience to guide circuit- and computation-level therapeutic strategies.

These results also indicate that key features of disease progression resemble the behavior of healthy individuals under high cognitive load. This suggests that, at least in early stages, the system is not operating in a fundamentally different mode, but instead closer to its limits. In healthy conditions, behavior is typically organized but can become disorganized under heavy load. In early ADRD, this transition appears to occur at lower levels of demand, reflecting a shift toward a less organized state. Consistent with this interpretation, our methylphenidate results suggest that aspects of this disrupted structure can be transiently restored.

### Longitudinal primate models link pathology to behavior

This work underscores the potential of longitudinal nonhuman primate models for studying early-stage neurodegenerative disease (Buffalo et al., 2019). Longitudinal monitoring of behavior and large-scale neuronal population activity in the same individuals makes it possible to track disease progression, identify early biomarkers, causally test hypotheses, and evaluate potential interventions in a controlled yet biologically relevant system. The similarity of monkey and human brains and behaviors enhances the translational potential of these approaches and findings. Collectively, these approaches enable investigation of intermediate population-level changes that link molecular pathology to complex behavior in ways that are difficult to achieve in other model systems.

### Conclusion and impact

Taken together, our findings suggest that early-stage ADRD progression is best understood as a progressive loss of organization that unfolds in parallel across levels of analysis (Figure 5). Despite preserved single-neuron tuning and intact encoding of many basic visual features, we observe coordinated changes in the structure of neuronal population activity, inter-area interactions, and visually guided behavior that accompany well-known changes in molecular and cellular structure. The parallel time course of these changes indicates that pathology does not simply eliminate representations, but disrupts the coordinated organization that enables distributed activity to support structured, goal-directed behavior. This perspective places disease progression along a continuum from cognitive resilience to cognitive disorganization and identifies organization as a property that links molecular pathology, population activity, network coordination, and behavior. In this view, neuronal population organization is a central substrate of cognitive integrity, and its degradation provides a mechanistic bridge between pathology and behavior.

---

**Supplementary Figure 1.**
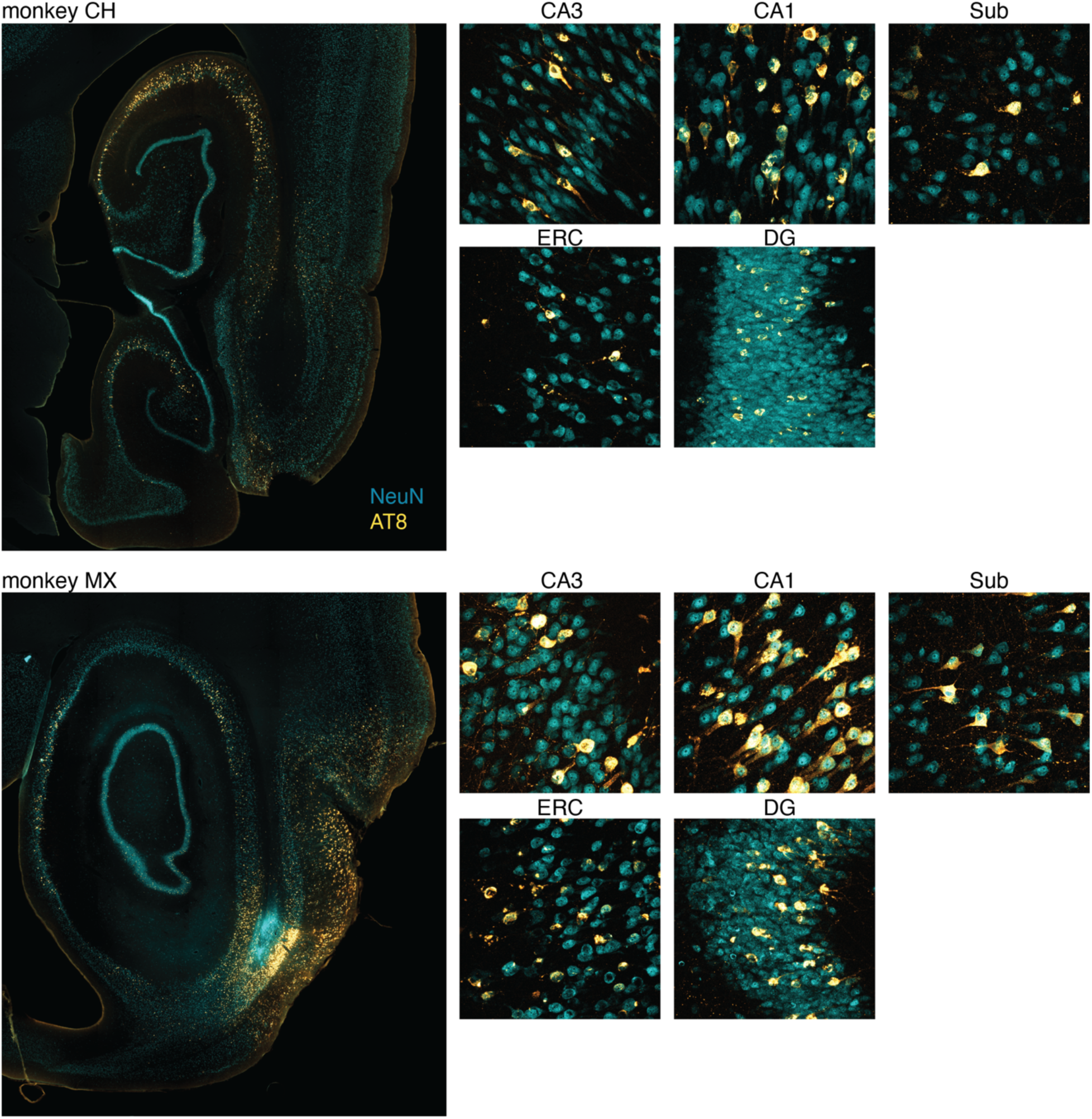
AAV-2xTau injection in the entorhinal cortex (ERC) induces robust tau propagation. Histology performed one year after viral injection of dual tau mutation in the ERC. AT8 (gold) was used as a marker for pathological tau spread, with NeuN (blue) labelling neuronal nuclei. No AT8 labelling would be expected in these animals in the absence of the AAV-2xTau injection (see (Beckman *et al*., 2021; Beckman *et al*., 2024)). Left hemisphere section shown for each animal. AAV injections were bilateral and tau was confirmed in both hemispheres. Increased magnification portions of the larger (left) image are shown for hippocampus (CA1 and CA3), subiculum (Sub), entorhinal cortex (ERC), and dentate gyrus (DG).

**Supplementary Figure 2.**
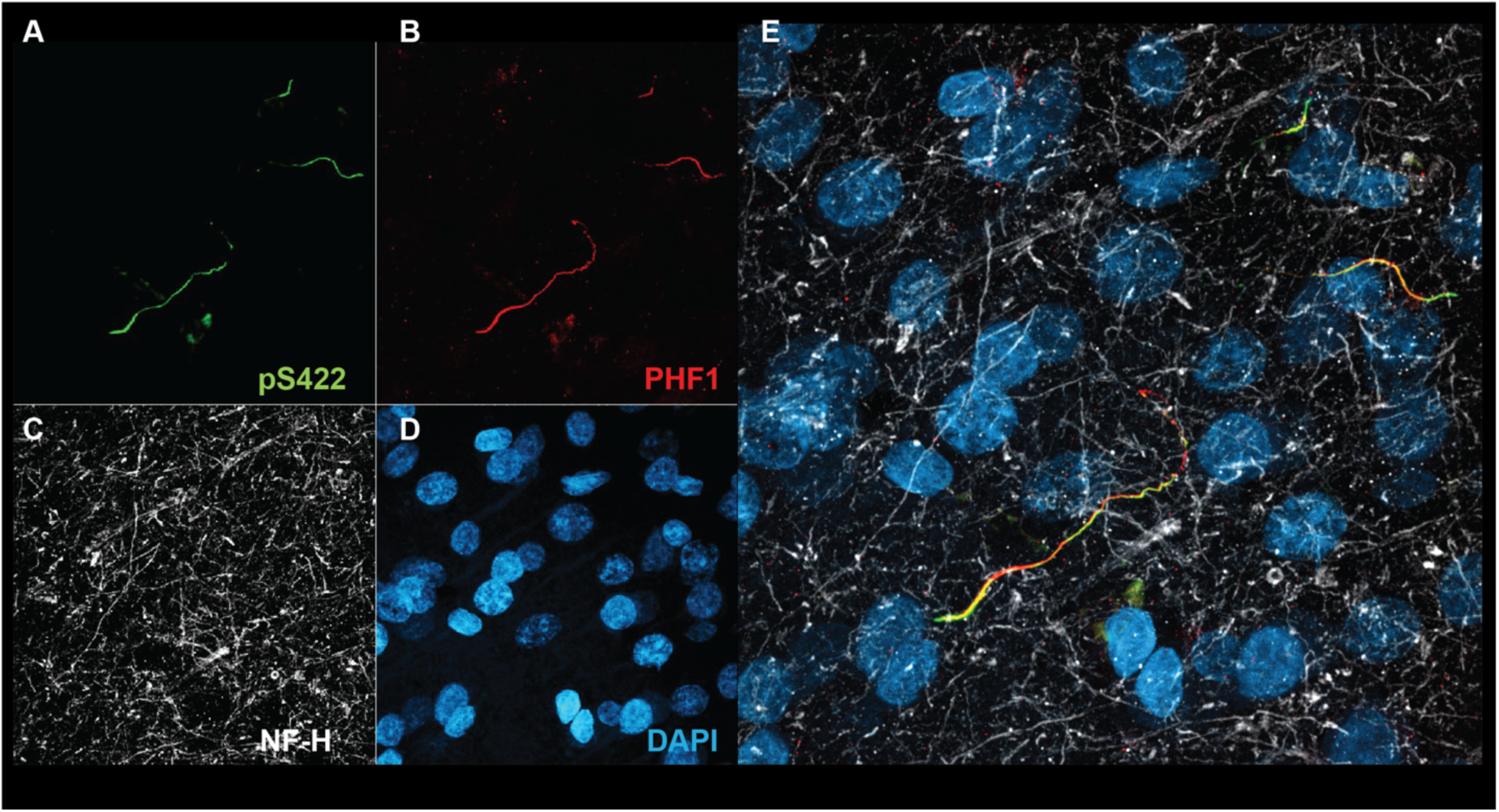
Elaboration of histological observations in V4. The same field of view from Figure 1C is shown by compartment for tau markers A) pS422 and B) PHF1, C) Neurofilament Heavy, NF-H, which labels neuronal cytoskeletal elements, particularly in axons, and D) DAPI, which labels nuclei. E) shows panels A-D combined.

## Methods

### Animals

Two adult male rhesus macaques (13 to 15 years old, conforming to the age ranges used in (Beckman *et al*., 2021; Beckman *et al*., 2024)) had bilateral stereotaxic injections of an adeno-associated virus expressing a human double tau mutation (AAV-P301L/S320F, 1.176 × 10^13 genomic copies/mL) into entorhinal cortex (ERC) (Beckman *et al*., 2021; Beckman *et al*., 2024). To increase the area of AAV expression, two injections were made per hemisphere. Injections were performed using the StealthStation surgical navigation system (Medtronic, Minneapolis, MN) using presurgical MRIs as guidance. During the same surgery, and following the AAV injections, we implanted microelectrode arrays (Blackrock Microsystems, 48 electrodes per array, arranged in a 6×8 rectangle) sub-durally in the left hemisphere of each animal. We implanted one array in parietal area 7a and two arrays in visual area V4. The arrays were connected to percutaneous connectors, which allowed electrophysiological recordings. The distance between adjacent electrodes was 400 μm, and each electrode was 1 mm long. All experiments were conducted with approval by the Institutional Animal Care and Use Committee (IACUC) at the University of California–Davis.

### AAV preparation

The adenovirus used in this work has been described in detail elsewhere (Beckman *et al*., 2021; Beckman *et al*., 2024; Koller et al., 2019). A capsid 1 adenovirus was packaged with one copy of human 0N/4R tau-containing two mutations (P301L/S320F) that render it more aggregation-prone than wild-type tau. The expression of this tau insertion is under the control of the hybrid cytomegalovirus enhancer/chicken β-actin (CMV/CBA) promoter, a CBA intron (first intron of chicken β-actin gene plus the splice acceptor of the rabbit β-globin gene), woodchuck hepatitis virus post-transcriptional regulatory element (WPRE), and bovine polyA. Virus packaging and purification were performed at the Penn Vector Core (University of Pennsylvania), following gold standard laboratory practices, including endotoxin testing (<5 endotoxin units per mL), purity testing (sodium dodecyl sulfate-polyacrylamide gel electrophoresis), and titration (three rounds of digital polymerase chain reaction). After preparation, aliquots of AAVs were stored at −80°C until surgery.

### Structural MRI and surgical planning

Pre-surgery structural imaging of both monkeys was performed using a 3T MRI scanner (Skyra, Siemens Healthcare, Germany) with an 8-channel receiver coil optimized for monkey brain scanning (Rapid MRI, Columbus, OH). T_1_-weighted images were acquired using the magnetization-prepared rapid acquisition with gradient echo (MPRAGE) pulse sequence with the following parameters: field of view = 154×154 mm; 480 sagittal slices; TR/TE = 2500/3.65 ms; flip angle = 7°; TI = 1100 ms; voxel size 0.6×0.6×0.6 mm, interpolated to a resolution of 0.3×0.3×0.3 mm. Anatomical images were used to identify the location of ERC, and, in conjunction with fiducial markers, determine appropriate injection sites.

### CSF, serum, and plasma collection and processing

Cerebrospinal fluid (CSF), plasma, and serum were collected before the initial surgery, approximately once per month post-surgery, and immediately before euthanasia. Before each collection, animals were sedated with ketamine (5–30 mg/kg, intramuscular) and dexmedetomidine (0.0075–0.015 mg/kg, intramuscular). To collect of CSF, a 23-gauge spinal needle was inserted into the subarachnoid space of the cisterna magna, and 1–2 mL of CSF was aspirated into sterile glass vials, which were then centrifuged at 400×*g* for 10 minutes at 4 °C to remove cellular debris. The supernatant was then aliquoted into low protein-binding cryotubes and stored at –80°C. To collect serum and plasma, ≈30 mL of venous blood was collected in EDTA-containing tubes (plasma only) and centrifuged at 1500×*g* for 15 min. The samples were divided into 0.5 mL aliquots and stored at −80°C.

### Phosphorylated tau quantification in biofluids

Plasma and CSF samples were analyzed using the NULISAseq^TM^ CNS Disease Panel 120 (Alamar Biosciences, Fremont, CA), a sequencing-based fluid proteomics platform with increased sensitivity and dynamic range compared to traditional quantification techniques (Feng *et al*., 2023). Briefly, specimens were assayed in singlets in the same plate to minimize batch effects. Each assay plate included multiple internal and external controls: three sample controls (derived from a well-characterized plasma source) and four negative control wells (buffer only, used to establish the limit of detection for each analyte), as well as internal controls (IC) spiked into every well to compensate for technical variability.

All assay steps were executed automatically using a preprogrammed ARGO HT system (Alamar Biosciences). Pooled sequencing libraries were generated and sequenced on an AVITI benchtop sequencer (Element Biosciences, San Diego, CA). Raw sequencing counts for each analyte were normalized to the corresponding IC counts and log2-transformed, yielding results in normalized protein quantification (NPQ) units. Scaling, normalization, and log2 transformation were performed automatically by the ARGO Control Center (ACC) software (Alamar Biosciences) based on the FASTA file generated by the sequencer.

Sample integrity was evaluated using three criteria: number of IC reads (> 1,000 reads), total number of reads (> 500,000), and sample IC reads relative to the plate median (+-40% of the plate median). All samples included in this study cleared all three criteria (IC reads min = 10,565; total reads min = 1,457,005 reads; IC% max = 22.56%). For the two targets reported in this study, pTau-181 and pTau-231, all samples were above the limit of detection. Intraplate coefficient of variation (CV%) was 11.37% for pTau-181 and 6.09% for pTau-231.

### Animal observation and health monitoring

The weights of both animals were tracked weekly and remained stable, with both slightly increasing across the year of testing. Both animals routinely received physical examinations as well as close observation of food intake and informal assessments of cage-side behavior. Throughout the year of testing post disease onset, both animals maintained stable relationships with their pair-housed partner and frequently engaged in routine social interactions and grooming bouts. Neither animal exhibited major behavioral changes that warranted external intervention. Further, throughout the year of testing, both animals remained willing to enter their chair and work for juice and food rewards in the experimental testing rig. The animal controlled the duration of each experimental session (a session ended when they lost interest in doing the task).

### Behavioral training and testing

During training and recording sessions, the monkeys sat facing a computer screen in a primate chair that provided access to a rigid straw that was used to deliver liquid rewards. During training and some testing sessions, padded spacers were placed near, but not touching, the sides of the head to encourage a forward-facing sitting position. In both animals, these were eventually abandoned during recording sessions. Eye tracking was used to control the behavioral task and successful tracking was possible when monkeys faced forward and placed their mouths on the reward straw. Otherwise, behavioral trials did not initiate. Monkeys were given *ad libitum* access to water in their home enclosure and were rewarded with juice and food rewards during training and testing. They received the remainder of their food rations after the completion of their daily testing.

The monkeys were trained for between 3 and 6 months prior to surgery to perform a variety of visually guided behavioral tasks. For all tasks, each trial began when the monkey fixated a small spot in a 1.0° circular fixation window in the center of a video display for 200-250ms. The color of the fixation spot was linked to the different tasks, but the monkeys only ever performed one task at a time, in blocks of trials that typically lasted for an entire experimental session.

This manuscript discusses three classes of behavioral tasks. The directed saccade (“gap”) task (Figure 1C), required the monkeys to maintain fixation on a central spot until a peripheral reward target appeared on the screen at one of four possible locations. After target onset, the monkey had to saccade to the target within 400ms to receive a liquid reward. During the initial fixation period, visual stimuli were displayed for either 200 or 300ms to measure response properties of the recorded visual neurons. A variety of stimuli were used, some of which are reported in Figure 3. Inspired by past studies using this task to investigate the ability to withdraw fixation and attend to peripheral targets (Abel *et al*., 2002), we varied the relative timing of fixation offset and target onset. The peripheral target could appear while the fixation spot was still on the screen, at the same time the fixation spot disappeared, or following the offset of the fixation spot. In all cases, the monkey was rewarded for making a saccade to the target within 400ms of its appearance on the screen.

In the rule-based visual foraging task (Figure 1D), the monkey fixated a central spot, and then an array of visual stimuli was displayed. After the offset of the fixation point, the monkeys were rewarded for fixating stimuli from a target category (e.g. animals) and not for fixating distractor stimuli (e.g. inanimate objects). Monkeys were rewarded the first time they fixated inside an invisible circle around a target stimulus and were not rewarded for repeatedly fixating the same target location or for fixating a distractor. The position of target and distractor stimuli changed on every trial and different target and distractor stimuli were used in different experimental sessions.

In the preferential looking task (Figure 1E; (Zola *et al*., 2013; Zola *et al*., 2000)), after the monkey fixated a central spot we displayed two visual stimuli in opposite hemifields. After 250ms, the fixation spot turned off. This indicated that the monkey could freely view the stimuli. The monkeys were rewarded for accumulating 1000 ms of total viewing time on either or both images. Visual stimuli were pseudo-randomly selected from the THINGS image set (Hebart et al., 2019) and the amount of visual exposure to each image was manipulated such that stimuli had either never been viewed before (novel) or had been previously viewed anywhere from once to many hundreds of times (familiar).

### Neuronal recordings

We recorded neuronal activity from the electrode arrays during daily experimental sessions for one year in each animal. Using our recording methods, it is nearly impossible to tell whether we recorded from the same single– or multi–unit clusters on subsequent days. To be conservative, the analyses presented here are performed on data from single recording sessions and then, when relevant, are compared across time.

An experimental session was included in neuronal analyses (Figures 3 and 4) if the monkey completed at least 35 behavioral trials (monkey CH, mean 62 trials; monkey MX, mean 45 trials) and a sufficient number of units responded at 10% more to the visual stimulus or target onset than when the monkey fixated a blank screen at the beginning of each trial. In Figure 4, the minimum was 45 active V4 units (monkey CH, mean 104 units, minimum 46 units, maximum 128 units; monkey MX, mean 75 units, minimum 46 units, maximum 122 units), 10 7a units (monkey CH, mean 22 units, minimum 11 units, maximum 38 units; monkey MX, mean 28 units, minimum 13 units, maximum 53 units), and 100 V4-7a unit pairs (monkey CH, mean 1382 pairs, minimum 115 pairs, maximum 4826 pairs; monkey MX, mean 1729 pairs, minimum 128 pairs, maximum 4270 pairs).

To ensure that changes in firing rates cannot explain the noise correlation changes reported in Figure 4 (Churchland et al., 2010; Cohen and Kohn, 2011; Ruff and Cohen, 2014), we calculated noise correlations by subsampling units such that there was a matched mean firing rate for all included units across the entire year of testing. Units were randomly dropped from inclusion until the distribution of baseline firing rates had an average between 20 and 30 spikes per second.

### Data analysis and statistical comparisons

We assessed the statistical significance of changes in behavioral and neuronal measures across time using F-tests on a linear model against the null hypothesis of a constant slope.

We assessed stimulus information in populations of neurons using linear regression with neurons as regressors, using leave one out cross validation across trials. We quantified decoding performance as the correlation between predicted and true stimulus or visual feature values.

### QR orthogonalization

To visualize changes in the structure of population activity (Figure 3B and 3D), we performed PCA on the average population responses to the stimuli shown in Figure 3A and then used linear regression to identify the best dimension for representing each of the two features of interest, color and shape. We next performed QR decomposition on these axes and projected responses into these orthogonalized dimensions. Responses were then scaled by subtracting the mean and dividing by the maximum response from each session so that responses could be compared across sessions.

### Intrinsic timescale calculation

To estimate the intrinsic timescale of the activity of each unit, we computed a shuffle corrected, mean-rate normalized, autocorrelogram (Smith and Kohn, 2008) using spike times during a 200 ms window of stable fixation on a blank screen at the beginning of each trial. We then fit an exponential decay function to each autocorrelogram and defined intrinsic timescale as the decay time constant. To account for refractory periods between spikes (Soyuhos *et al*., 2025), we began the fitting procedure 3ms after the 0 lag. We included units that were well fit by an exponential function (those with goodness of fit (R^2^) > 0.5). This analysis was only performed on V4 data because of an insufficient number of units in 7a whose autocorrelograms were well fit by an exponential function.

### Behavioral entropy calculation

To quantify the structure of visual foraging behavior (Figure 4E, F), we computed a transition entropy metric based on sequences of fixations within each trial. For each fixation location, we calculated the proportion of next fixations at each of the other locations. We defined transition entropy as the Shannon entropy of this distribution,

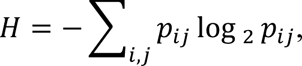

where *p_ij_* denotes the probability of transitioning from location *i* to location *j*. Higher entropy values indicate more uniform, less structured transition patterns, whereas lower entropy reflects more stereotyped or organized foraging behavior. To normalize for factors related the number of fixations within each trial, we generated a null distribution by randomly shuffling the order of fixations within each trial and recomputing entropy. Entropy values were z-scored relative to this shuffled distribution (*z* = (*H*_actual_ − μ_shuffled_)/σ_shuffled_), such that negative values indicate more structured behavior than expected by chance, and values near zero indicate approximately random foraging. This metric captures the degree of structure in fixation sequences without imposing assumptions about specific search strategies or saccade statistics. We included sessions in this analysis if there were at least 20 successfully completed trials.

### Convolutional neuronal network model

To generate hypotheses about how disease progression might affect stimulus representations, we used activations from a pre-trained convolutional neural network (VGG-16). We measured model responses to our stimuli and used PCA to reduce the dimensionality of responses from the pooling layer neurons of model layer 4. We then decoded stimulus features (Figure 3E) while randomly removing different numbers of dimensions to simulate disease related changes.

### Methylphenidate administration

Methylphenidate was orally administered for short bouts starting approximately six months after disease onset. We used a dosage of 3.5 mg/kg based on our previous studies (Ni *et al*., 2022). In monkey CH, the drug was administered on 18 days during three-plus weeks of testing starting on day 176, while in monkey MX it was given on 5 days during one week of testing starting on day 174. The methylphenidate was dissolved in 20-30 mL of flavored sugar water and administered orally using a method adapted from (Ni *et al*., 2022; Soto et al., 2012). As in our previous study, methylphenidate was administered 30-45 min before the start of the experiment (Ni *et al*., 2022). An additional 6 day bout of MPH was administered in monkey CH starting on day 251, but an insufficient amount of data were collected during the time spans to expand the fitting performed in Figure 4E.

### Perfusion and tissue preparation

Necropsy was performed by a team of trained pathologists in a biosafety level 2 environment. On the day of euthanasia, monkeys were deeply anesthetized with ketamine (25 mg/kg) and transcardially perfused with saline until the vascular bed was cleared, as described previously (Beckman *et al*., 2021; Beckman *et al*., 2024). After perfusion, brains were harvested, hemisected, and sectioned into 6 mm-thick coronal blocks. Blocks were fixed by immersion in 4% paraformaldehyde with 0.125% glutaraldehyde in phosphate buffer pH 7.4 at 4°C for 48 h under agitation. After the washes, blocks were stored in phosphate-buffered saline (PBS) with 0.1% sodium azide at 4°C until processed. Semiseriated, 30 μm-thick coronal sections were obtained by cutting blocks containing the ERC and HF in a freezing microtome. Sections were stored in antifreeze solution with 0.1% sodium azide at -20°C.

### Immunofluorescence and histological staining

We used immunofluorescence to validate the AAV injection sites and expression of tau proteins, based on the protocol in our previous studies (Beckman *et al*., 2021; Beckman *et al*., 2024). Nuclei were stained with 4′,6-diamidino-2-phenylindole (DAPI 0.001 mg/mL; Invitrogen). Neuronal markers NEUN (1:1000; 266 004, Synaptic Systems GmbH, Göttingen, Germany) and Pan-Neurofilament (panNF; 1:400; DMAB 7133, Creative Diagnostics, New York, NY) and tau epitopes AT8 (1:500; MN1020, Invitrogen, Waltham, MA), pS422 (1:1000; ab79415, Abcam), and PHF1 (1:1000, generously provided by Dr. Nicholas Kanaan) were used for validation.

### Microscopy, image analysis, and histological validation

All images were acquired using an upright microscope (AxioImager Z2; Carl Zeiss AG, Oberkochen, Germany) equipped with an Axiocam 620 mono camera, LSM800 confocal head, and a 32-channel Airyscan detector (Zeiss). Low-magnification tiles were acquired using single-plane widefield illumination, while high-magnification images were acquired as 3D stacks (20-30μm) and collapsed via maximum-intensity projection for reporting. Images were acquired using the ZEN core software (Zeiss). Brightness and contrast were adjusted in uniform manner across images. To ensure that the model was successfully induced, we identified injection tracks pointing to the entorhinal cortex (ERC), and markers of tau.

## Acknowledgements

Technical assistance: Lisa Novik, Stephanie Hawbecker, Sarah Grisso, Chloe Meier, Naomi Richards, Laura Garzel, Kursti Eaves.

## Authors contributions

D.A.R., M.R.C., and J.H.M. conceptualized the experiment. D.A.R. and M.R.C. designed the experiments and wrote the paper. D.A.R. performed the experiments. D.A.R., D.E.G.S., and R.S. performed data analysis and modeling. G.B.D., D.J.G., D.B., S.O., K.S., and C.E. performed tissue and biomarker collection and analysis and histological processing and imaging. D.A.R, M.R.C., S.O., S.M., and J.H.K. performed surgical planning and execution. J.H.M. and M.R.C. supervised and funded the work. All authors provided comments and discussion about the manuscript.

## Data, code, and materials availability

Data deposited in repository upon publication

## Supplementary Materials

Materials and Methods

Figs. S1 and S2

